# Managing mixed stands can mitigate severe climate change impacts on ecosystem functioning

**DOI:** 10.1101/2020.06.13.149856

**Authors:** M. Jourdan, T. Cordonnier, P. Dreyfus, C. Riond, F. de Coligny, X. Morin

**Affiliations:** CEFE UMR 5175, CNRS - Université de Montpellier - Université Paul-Valéry Montpellier - EPHE, 1919 Route de Mende, F-34293 Montpellier Cedex 5, France; Agence de l’environnement et de la maitrise de l’énergie, ADEME, Angers, France; ESE, CNRS, AgroParisTech, Univ. Paris-Saclay, F-91400 Orsay, France; Université Grenoble Alpes, INRAE, UR LESSEM, 38000 Grenoble, France; INRA, UR 0629 URFM Ecologie des Forêts Méditerranéennes. Centre de Recherche PACA, Avignon, France; ONF, Office National des Forêts - Département RDI, Avignon, France; ONF, Office National des Forêts - Département RDI, Chambéry, France; INRAE, UMR931 AMAP, Botany and Computational Plant Architecture, TA A-51/PS2, Boulevard de la Lironde, 34398 Montpellier Cedex 5, France

**Keywords:** Species diversity, mixed forests, mountain forests, gap-model, management, climate change

## Abstract

Climate change affects forest ecosystem processes and related services due to increasing temperature and increasing extreme drought event frequency. This effect can be direct through the alteration of the physiological responses of trees, but also indirect, by modifying interactions between trees and thus changing communities’ composition. Such changes might affect species richness with high impacts on ecosystem functioning, especially productivity.

Regarding management issues, mixed stands are usually considered a good option to maintain forest cover and ecosystem services under climate change. However, the possibility to maintain these mixed stands with management actions with positive effects on forest functioning under climate change remains uncertain and deserves further investigations. Relying on a simulation-based study with a forest gap model, we thus addressed the following questions: (1) Are monospecific stands vulnerable to climate change? (2) Would mixed stands significantly mitigate climate change effects on forest productivity and wood production under climate change? (3) Would conversion to mixed stand management affect significantly forest productivity and wood production under climate change compare to monospecific management?

With a 150 years simulation approach, we quantified potential climate change effect (using RCP 8.5) compared to present climate and managements effect in the French Alps, focusing on five tree species. The gap-model we used included a management module, which allowed testing six silvicultural scenarios on different stands, with various composition, structure or environmental conditions, under climate change.

These simulations showed that monospecific stands currently growing in stressful conditions would be too vulnerable to climate change to be maintained. Managing mixed stands or conversion from pure to mixed stands would make it possible to maintain higher productivity in the long-term than monospecific stands, even under severe climate change. This pattern depends to species and sites considered. Our results will feed into discussion on forest management in the context of climate change.

## 1 INTRODUCTION

In the Northern hemisphere, climate change will lead to increased temperature and changes in precipitation regime (not spatially homogeneous) as well as more frequent and more intense extreme climatic events during the next decades (Pachauri et al., 2014), particularly with strong drought and/or thermal stresses. Such changes, and notably extreme drought events, may be very damaging for European ecosystems (Maracchi et al., 2005) and especially for forests. Warmer and drier conditions can lead to medium or long-term damage (Bréda et al., 2006a; Linares et al., 2010), including massive mortality events. Thus longer, more intense and more frequent droughts may induce forest stands dieback, as already reported (Allen et al., 2010; Bigler et al., 2006a; Guarín and Taylor, 2005), sometimes with a time lag of several years (Bigler et al., 2007). Furthermore, beyond these direct impacts of climate change on tree physiology, climate change effect can be also indirect by altering communities composition and species richness (Bertrand et al., 2011; Lenoir et al., 2008), which in turn will impact ecosystem functioning (Loreau, 1998), and stressful climatic events can also increase the vulnerability of forest stands to pathogen attacks (Desprez-Loustau et al., 2006) or fire risks (Dale et al., 2001).

Numerous studies have shown that species richness may strongly modify ecosystem functioning, and especially increase productivity (Cardinale et al., 2007; Hooper et al. 2005). Focusing on forests, a few experimental (Jones, McNamara, & Mason, 2005; Pretzsch, 2005), observation-based (Forrester et al., 2016; Liang et al 2016; Toïgo et al., 2015) and modelling studies (Morin et al., 2018, 2011) have also found that diversity may lead to an overyielding effect in comparison with monospecific stands. Several studies focusing on diversity-productivity relationship have also aimed at exploring the impact of climate on these patterns (Jactel et al 2018, Blois et al., 2013; Paquette and Messier, 2011). Due to these positive effects, favoring species richness or mixed stands has been considered a good option to mitigate climate change negative impacts on forest ecosystem functioning (e.g. Hisano et al., 2018).

Recent studies have shown that even only two species stands can mitigate climate change impacts, through increasing and/or stabilizing stand productivity (Del Río et al., 2017). Thus, mixed stands management could be an efficient solution to sustain forest functioning and better preserve their services (e.g. Schwaiger et al., 2018). However, results are actually more contrasted when considering other kinds of mixed stands, with either positive or negative effects of species mixing on ecosystem functioning (Grossiord et al., 2014; Merlin et al., 2015; Jourdan et al. 2019), depending on environmental conditions and species composition (Grossiord et al., 2014; Jucker et al., 2014). For instance, mixed stands can have a negative effect on stand response to drought stress (Grossiord et al., 2014b). Better understanding how mixed stands may behave differently from monospecific stands in various ecological conditions is thus required, for instance by using pseudo-experimental approaches on specific mixed stands (del Río et al., 2014; Jourdan et al., 2019; Pretzsch et al., 2013) or experimental approaches (e.g. BIOTREE (Scherer-Lorenzen et al. 2007) or ORPHEE (Castagneyrol et al 2014)). In the first case, climate change effect is taken into account indirectly according to the space-for-time substitution, using environmental gradients (Jourdan et al., 2019; Jucker et al., 2014a), for instance latitudinal or altitudinal gradient. In the second case, the number of tested forest types is limited, because such experimental protocols are difficult to carry-out. In Europe, the majority of temperate forests are managed (Morneau et al., 2008) and the main stand structural characteristics (total basal area, tree density, species composition) of these forests have been controlled since up to several centuries (Reineke, 1933). In a context of energy transition, in which wood resource is more and more targeted by public policies, developing sustainable forest management under climate change that fulfills both ecological and economic challenges appears an essential task to maintain every forest ecosystem services. Stand management can be an efficient tool to mitigate the negative impacts of intensive droughts and promoting forest adaptation (Millar et al., 2007), through controlling density (Trouvé et al., 2017) or structural heterogeneity (Cordonnier et al., 2018a) including species mixing. It is however difficult to anticipate the combined effect of climate change and management on the long-term maintenance of species mixing in forest stands as well as the resulting effect on forest functioning (Cordonnier et al., 2018).

Yet, quantifying and predicting ecosystem change in composition and functioning under climate change and with forest management remains a difficult task (Morin et al., 2018), because of the great uncertainty in climate changes prediction (IPCC – Pachauri et al., 2014- proposes several climate scenarios) and the difficulties to forecast synergy between climate change and its direct and indirect impacts. In context embedding so many uncertainties, modelling approaches may be pivotal to test climate change impacts and management effects on forest ecosystems (Ameztegui et al., 2017; Reyer et al., 2015). Forest models integrating climate allow to consider climatic variation effect in their predictions of future forest structure and ecosystem services delivery (for example gap-model: ForClim, Bugmann, 1996), contrary to other models which are calibrated with past dendrochronological data without considering climate effects (relevant only in a constant and not dynamic climate, eg. Fagacée Le Moguédec and Dhôte, 2012). Models simulating also forest dynamics on forest massif scale and calibrated on Alps are few (ForClim, Bugmann, 1996, ForCEEPS, Morin et al., 2020b). Models allow working on long-term time scales (up to several hundred years) and simulating the impact of future climate change (i.e. not like dendrochronology studies that focus on the impact of past climate). Then this kind of tools is especially relevant to explore the effect of different stand management, whereas it is more difficult with other type of model or with experimental approaches in forests.

In this study we used the forest dynamics gap-model ForCEEPS to carry-out forest simulations in the French Alps integrating climate change and forest management and the five most common species in these forests. We simulated monospecific and mixed forests (some combinations of the five species) over the next century in four sites in French Alps. The simulations were run using a severe climate change scenario and by applying six different management scenarios to optimize wood production (some of them being oriented towards the promotion of mixed stands). More specifically, we aimed at answering the following questions (see also Figure S1):

1. Are monospecific stands vulnerable to climate change?
2. Would mixed stands significantly mitigate climate change effects on forest productivity and wood production under climate change?
3. Would conversion to mixed stand management affect significantly forest productivity and wood production under climate change compare to monospecific management?

## 2 MATERIAL AND METHODS

### 2.1 Description of the forest dynamics model

#### 2.1.1 General description

ForCEEPS is a forest dynamics model simulating population dynamic of one or several tree species in small parcels of land (“patches”) (Morin et al., 2020b). The model is individual-based, and predicts forest composition, biomass, and productivity, by considering abiotic (climate and soil properties) and biotic constraints (competition for light) to tree establishment, growth, and survival. The patches are independent, and dynamics at the forest level are obtained by aggregating patches together (Bugmann, 2001).

The main processes included in the version of ForCEEPS used in the present study are derived from FORCLIM 2.9.6. (Bugmann, 1996), except that the dynamics are simulated at the individual level (and not at the cohort level).

Tree establishment is determined by species-specific responses to five factors: minimum winter temperature, degree-days sum during the growing season, soil water content, light availability, and browsing pressure (Bugmann, 1996). The model does not consider seed and seedling stages; thus, saplings are established with a diameter at breast height of 1,27 cm. Dispersal limitation is not taken into account in this model and patches receive an annual seed rain of all species included in the simulation, assuming the presence of seed-bearing trees around the simulated forest (Bugmann, 2001).

Tree growth (high, diameter, and other) depends on a species-specific optimal growth rate that is modified by abiotic (temperature, soil water content – SWC -, and nitrogen content in the soil) and biotic factors (size-dependent competition between trees). The main mechanism driving interactions among trees is competition for light. Forest successional dynamics is triggered by canopy-gaps, and thus relies on differential species growth responses to light conditions. Species with different shade tolerances have different light response curves. In full light, light-demanding species (i.e. mainly early successional species) grow faster than shade-tolerant species (i.e. usually late-successional species) that have, on average, a weaker maximum growth rate. As light availability decreases, the realized growth rate of shade-tolerant species becomes relatively stronger than shade-intolerant ones because of decreasing light availability. Furthermore, although each species has species-specific tolerances to environmental drivers, there is no competition for SWC and soil nitrogen taken into account in the model. However, SWC varies across years depending on temperature and precipitations of the site.

Tree mortality is driven by both stochastic and deterministic processes and depends on two components: (i) a ‘background’ mortality, and (ii) a stress-dependent mortality. Background mortality is a stochastic process occurring at low frequency, increasing with trees age and depending on species’ maximum longevity. Stress-dependent mortality relies on the growth pattern of each tree: if a tree grows very slowly during several successive years, it is more prone to die than a tree with a better growth. ForCLIM, from which ForCEEPS is derived, is a well-established model that has been validated by showing its ability to reproduce observed vegetation patterns in central European forests through a range of climatic and environmental conditions (see Bugmann, 1996, Didion et al., 2009). Recently, ForCEEPS was thoroughly calibrated and validated for the main forest types in French territory (Morin et al., 2020b), including French Alps, showing a strong ability to reproduce potential species composition, and stand productivity under current climate. With a newly developed management module, it appears as a robust tool to test composition and management effect on Alps forests.

The model works with climate time-series and can thus consider either data reflecting current climate or future climate scenarios. ForCEEPS now includes a new silvicultural module able to simulate successive thinning operations, as described in Appendix S2. We defined and used silvicultural scenarios that differed according to the targeted basal area after each thinning, rotation (time between each thinning), and targeted proportion of each species. At each thinning, trees are logged until objective basal area is reached, with from above or from below thinning.

#### 2.1.2 Species parameters

In ForCEEPS, each species is defined by 13 parameters that have been estimated from a large body of literature data and experimental or observational data. Hereafter we consider these parameters as species ‘traits' as they determine physiological species responses to environmental conditions. The variability among traits reflects several trade-offs of tree life-history strategies. It is worth noticing that observed functional patterns at community scale are emergent properties from species responses to processes embedded in the model.

#### 2.1.3 Sites and species

##### Studied sites

Simulations ran on four sites in the French Alps, covering a large latitudinal gradient related to important variations of temperature and precipitation: Bauges, Vercors Méaudre, Vercors Lente and Mont Ventoux (Table S3). As requested in ForCEEPS, each site was characterized by latitude, soil field capacity and slope. The climate of each site was characterized by monthly mean temperatures and monthly sums of precipitation, with inter-annual variations. Two different elevations were tested for each site, one at 1000 m (low elevation) and 1300 m (high elevation).

##### Studied species

We considered five species in this study: common beech *(Fagus sylvatica),* spruce *(Picea abies),* silver fir *(Abies alba),* Scots pine *(Pinus sylvestris)* and pubescent oak *(Quercus pubescens).* These species are widespread in the French Alps and represent economic issues (e.g. fir and spruce) or patrimonial interest in mountain forest (e.g. beech).

These five species allowed studying various types of mixtures, because of the physiological and ecological differences between species. Beech and oak are broadleaved species while spruce, fir and Scots pine are coniferous species. Beech and fir are late-successional and shade tolerant species, while spruce is a mid-seral species. Spruce is very sensitive to high temperatures in summer, and is the most drought-sensitive species (Caudullo et al., 2016). Beech is also sensitive to water stress, but recovers easily after an intense drought event (Lebourgeois et al., 2005). Silver fir is less sensitive to drought, but grows better in humid conditions (Lebourgeois et al., 2010; Mauri et al., 2016) while Scots pine and pubescent oak are more early succession and light-demanding species, tolerating drier conditions (Pasta et al., 2016).

Even if we considered just five species at the beginning of the simulations, we allowed other species to colonize the patches during the simulations due to their observed abundance in the sites (sycamore maple – *Acer pseudoplatanus* – or mountain ash tree – *Sorbus aucuparia).*

#### 2.1.4 Climatic data

In this study, we looked for comparing simulation results between current climate and changing climate. Climate variables were delivered by Météo France (available in «Drias», Météo-France and project GICC Drias – CERFACS, IPSL). For the simulations under current climate, we used monthly temperatures and precipitation for the last 50 years. Then we created climate series, selecting randomly yearly climate conditions (monthly temperature and precipitation), to obtain climate 150 years’ time-series of “stable” conditions between 2000 and 2150, i.e. with inter-annual variability but without any long-term trend in variable means. For the simulations under climate change, we used data generated according to the RCP 8.5 scenario (IPCC – Pachauri et al., 2014), i.e. the most extreme available scenario (in temperature and precipitation) as our aim was to address the potential mitigating effect of managed mixed stands under sever climate change. Climate data for the future are available between 2000 and 2100. For our simulations, we needed to work on 150-years period because of the timing of forest dynamics. Thus, we generated climate data between 2100 and 2150 by selecting randomly yearly climate conditions (monthly temperature and precipitation) between 2075 and 2100, i.e. stabilized conditions after 2100.

#### 2.1.5 Simulation design

In these analyses we used six silvicultural scenarios. They differed according to the basal area *(BA)* remaining in the stand after thinning (20m^2^/ha, 30m^2^/ha, or 80% of *BA* before thinning) and objective composition (“stable management” and “conversion management”). “Stable management” was applied to monospecific and mixed stands, according to which thinning operations aimed at conserving initial species composition. “Conversion management” consisted in thinning operations that preserved not-aimed species – up to a stated abundance – in a monospecific stand. Regarding mixed stands, the targeted distribution of relative abundances in the “stable” option are 50-50 or 80-20 for two species stands and 40-40-20 for three species stands. In the “conversion” option, the targeted distribution of relative abundances was 50-50 for two species mixed stands, and 30-30-40 for three species mixed stands (with 40% of the initial species of monospecific stand). In each case, rotation time was 12 years, as recommended by silvicultural guides of French northern Alps (Gauquelin & Courbaud, 2006). Each configuration of forest composition is shown in Figure 2. We used three different simulation designs to answer to our three questions (represented by colors).

**Figure 1:**
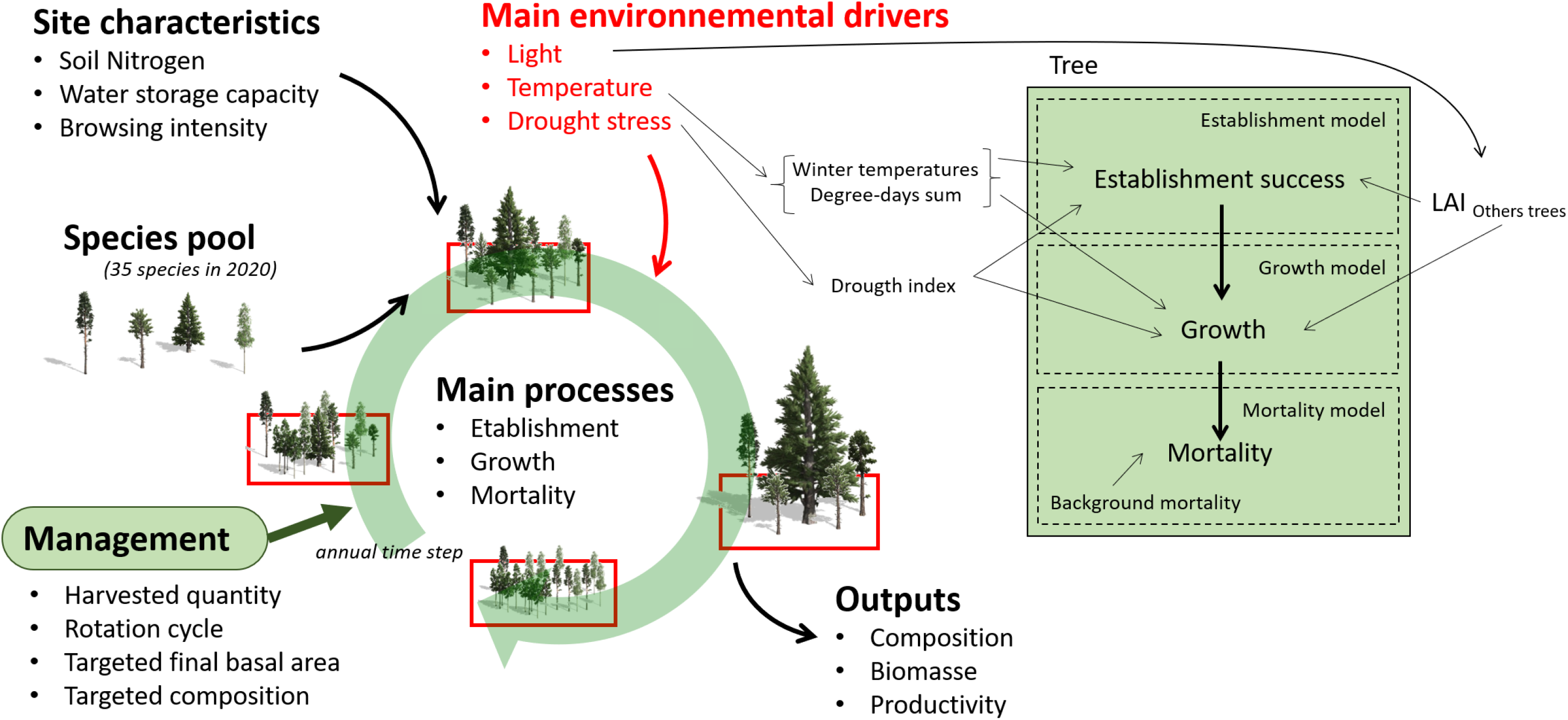
Simplified scheme of the gap-model ForCEEPS. The cycle, represented by the green arrow, represents the forest dynamics observable at patch scale, with recruitment, growth and mortality. The central diagram represents the variation of the basal area of a patch during a simulation.

**Figure 2:**
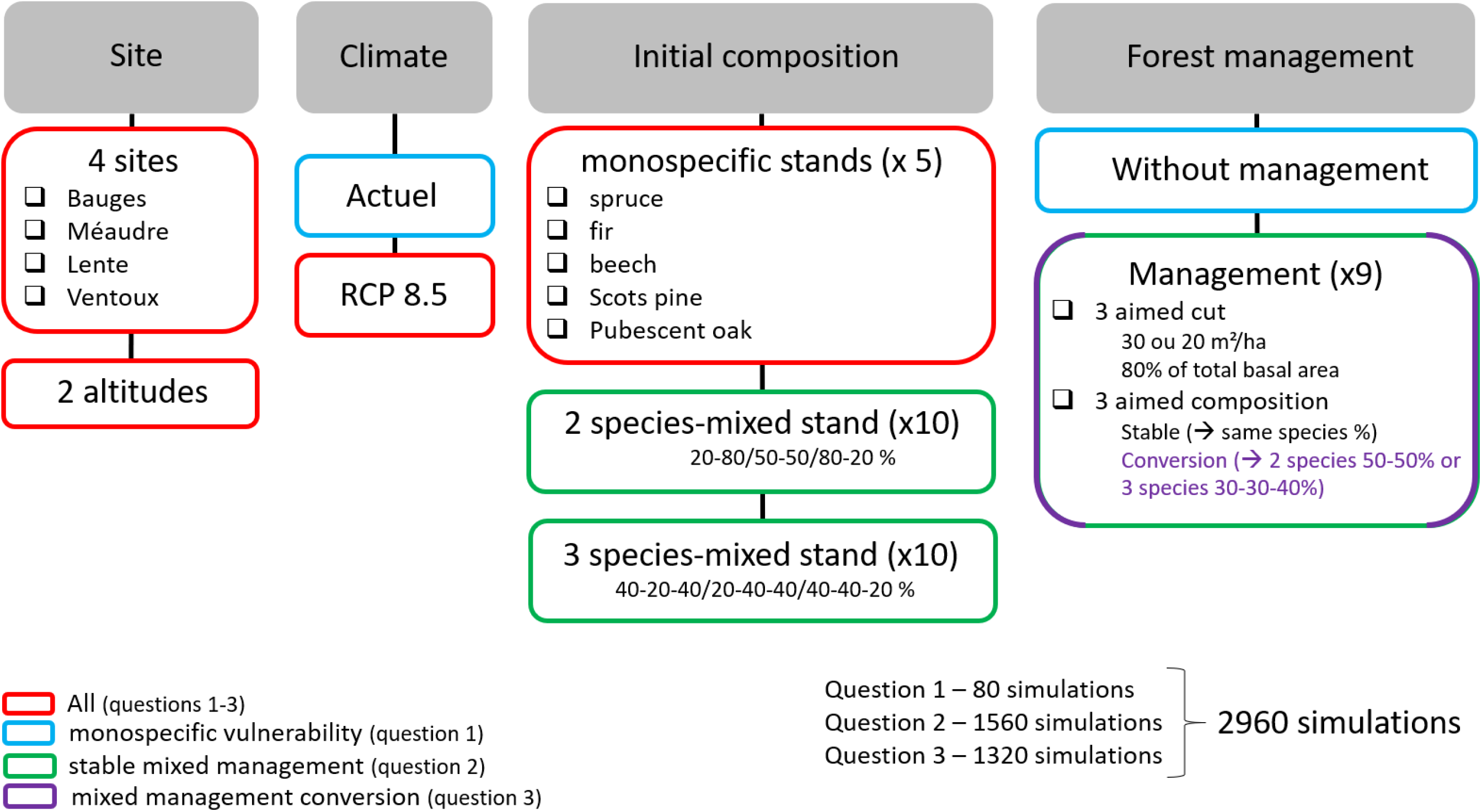
Scheme of the simulation design used for each question mentioned in the introduction: (1), represented by red and blue; (2), represented by red and green; (3), represented by purple and red.

First, to study monospecific stands vulnerability to climate change (Question 1), we worked on the four sites, considering both altitudes and with both climate scenarios (current climate and climate RCP 8.5). The simulations started from mature forest inventory, representative of Alps forest.

Note the the other species were allowed to colonize the site over the simulation. In this part, we did not consider any management actions, because we aimed at assessing the intrinsic stand vulnerability to climate change.

Then for studying mixed management effect on stand productivity and maintenance (question 2 and 3), we also worked on the four sites and considering both altitudes, but for climatic conditions we focused on RCP 8.5 climate. We considered monospecific and 2- and 3-species mixed stands. We worked with “stable management” and “conversion management” options for each of the three defined thinning objectives (see above). Note that the other species were allowed to colonize the site over the simulation. Figure 2 summarizes simulation plan of this study.

Preliminary analyses showed that simulations with the same characteristics (site, initial inventory, and climate) and only varying for their stochastic parameters used in some processes (mortality, recruitment) led to similar results (in terms of productivity, biomass, and composition). We thus decided to perform only one simulation per case, i.e. one initial inventory for one altitude/site/climate/management, for the sake of simulation time. Each monospecific and mixed stand simulations was simulated at each site and altitude on 50 independent patches of 1000 m^2^ (i.e. 5 ha in total). To study a complete stand rotation, simulations were run over 150 years.

### 2.2 Analyses

#### 2.2.1 Vulnerability of monospecific stands

We studied monospecific stand vulnerability through three indices. We considered final stand composition (calculated with basal area proportion of each species at the end of the simulation, i.e. after 150 years) as the stand may experience the colonization by the other species during the simulation. We also studied final basal area as a proxy of forest cover. We considered monospecific stands as stands with more than 70% (on basal area) of the targeted species.

##### Species vulnerability

To determine vulnerability of each monospecific stand, we needed to compare final proportion of the target species, between current and future climate conditions. We quantified species vulnerability to climate change by final proportion in the stand proportion of targeted species between current and future climate conditions.

##### Vulnerability of mean productivity

Mean annual productivity (in *BAI,* i.e. m2/ha/an) was calculated over three periods of 50 years: from year 1 to 50 from year 51 to 100, and from year 101 to 150. Working on three time periods allowed assessing variations in stand properties in the short and middle term. The vulnerability to climate change was assessed by comparing monospecific stand productivity under current and future climate conditions. Monospecific stands vulnerability to climate change is quantified by stand productivity proportion between climate change and current climate. We considered each monospecific stand separately for each site and each period.

##### Vulnerability of stand basal area

The vulnerability to climate change was assessed by comparing stand final basal area, i.e. proxy of biomass after 150 years of simulation, under current and future climate conditions. We considered each monospecific stand separately for each site and each period.

#### 2.2.2 Mixed *vs.* monospecific stand management

We compared mixed (2 or 3-species) and monospecific stand management using wood harvested (m2/ha) by timbering. We first carried-out this analysis for the “stable management”. Then we compared several “conversion management” scenarios using wood harvested (m2/ha) by timbering. Aim is to evaluate management effect on initially monospecific stand. To assess the effect of management, the comparisons were tested a t-test.

## 3 RESULTS

### 3.1 Are monospecific stands vulnerable to climate change?

#### 3.1.1 Stand vulnerability related to species identity

##### Under current climate

After 150 years of simulation under current climate, the relative abundance of the main species in monospecific stands may decrease significantly, depending on the species and the site considered (Fig. 3). Monospecific Scots pine stands did not persist on any site under current climate: less than 25% for Scots pine in Vercors (Fig. 3-B and C) and Bauges (Fig. 3-A), substituted by spruce, fir and beech. Monospecific spruce stands were not grown in Ventoux (Fig. 3-D) under current climate (less than 25%, mostly substituted by beech and fir), but showed a better ability to remain monospecific stands in Bauges and Vercors (close to 70%, Fig. 3-A, B and C). Monospecific fir and beech stands showed also a strong ability to remain monospecific stands over time, with a decrease in relative abundance between 30 % and 40%. Monospecific oak stands persisted in current climate in Bauges and Ventoux, with a decrease in relative abundance around 40%, and in Vercors (depending on elevation).

**Figure 3:**
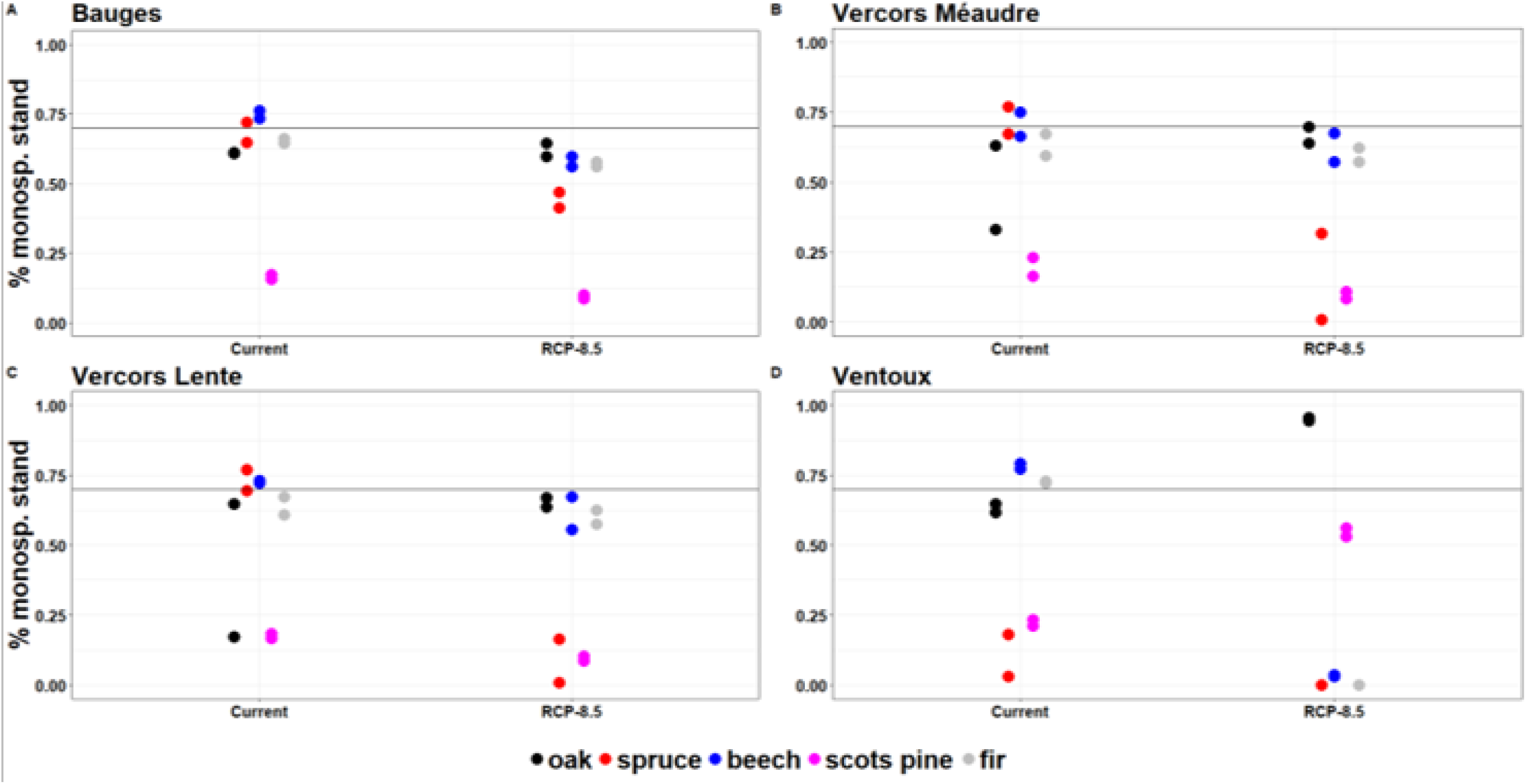
Final proportion of the target species in monospecific stands without management for each set of climate conditions: current climate or climate change (RCP 8.5). Each species is represented separately: beech (blue), Scots pine (pink), fir (grey), spruce (red) and pubescent oak (black). Each panel refers to one site: Bauges (A), Vercors Méaudre (B), Vercors Lente (C) and Ventoux (D). The black line represents the chosen threshold between monospecific (proportion higher than 70%) and mixed stand (proportion lower than 70%).

##### Under climate change

After 150 years of simulation, we found that climate change greatly impacted monospecific stands in each site, with an intensity depending on species and site considered. For monospecific spruce stands, the results showed a higher loss in spruce proportion for all sites, until total spruce disappearance (Ventoux and Vercors Lente). Monospecific spruce stands were very vulnerable in Vercors sites, with a predicted final proportion between 0 and 35% (Fig. 3-B and C). The vulnerability was lower in Bauges (Fig. 3-A, spruce stands final proportion remain higher than 40%). For monospecific Scots pine stands, the proportion remained comparable under climate change (around 20%), except in Ventoux (Fig. 3-D), where monospecific stands were much better maintained (more than 50% instead of 20%). For monospecific fir and beech stands, climate change effect depended on the site: proportion at the end of the simulation decreased in Ventoux (Fig. 3D: from 75% to 0%) substituted by Scots pines and other species, and slightly decreased in Vercors (Fig. 3-B and C) and Bauges (Fig. 3-A). Monospecific beech and fir stands were highly vulnerable in Ventoux and benefited from the strong vulnerability of spruce in Vercors sites. Oaks proportion in initially monospecific oak stands remained similar in climate change (compared to current climate) in Bauges and became higher in average in Vercors (from less than 50% to 70%) and in Ventoux (from 70% to more than 90%). Other species represented less than 10% of final *BA*, regardless of site and species, except in Ventoux.

##### Synthesis

Monospecific spruce stands in Ventoux and monospecific Scots pine stands in Vercors and Bauges did not grow at all (under current climate and climate change). Monospecific spruce stands were very vulnerable in Vercors under climate change. Monospecific beech and fir stands were very vulnerable in Ventoux. According our results, pubescent oak was less vulnerable under climate change compared to under current climate, in almost every case. Vulnerability remained limited in other cases.

#### 3.1.2 Stand vulnerability through average productivity

In Bauges and Vercors (Fig. 4), monospecific stands productivity were not affected by climate change. In Ventoux (Fig. 4), monospecific stands were vulnerable for all species at middle-term (51-100 years), but not at long-term (101-150 years). Between 51 and 100 years, species productivity decreased drastically. Then, between 101 and 150 years, productivity recovered at equivalent or greater level than monospecific stands in current climate, except for oak stand (with productivity almost divided by two). Because of change in species composition, climate change effect did not affect mean productivity of monospecific stands in the long-term.

**Figure 4:**
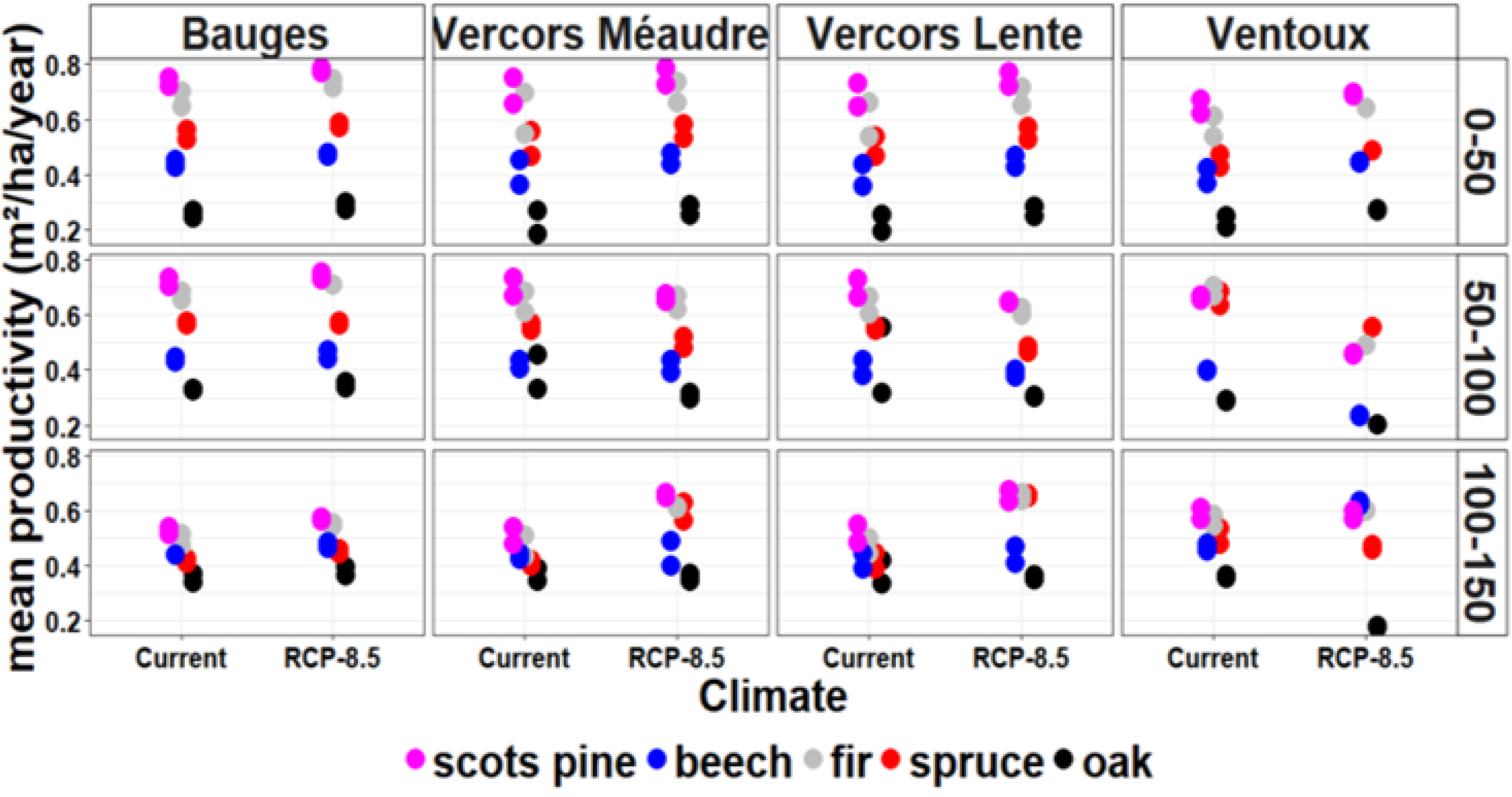
Mean productivity of the target species in monospecific stands without management for each set of climate conditions: current climate or climate change (RCP 8.5). Each species is represented separately: beech (blue), Scots pine (pink), fir (grey), spruce (red) and pubescent oak (black). Each column represents each site, from left to right: Bauges, Ventoux, Vercors Lente and Vercors Méaudre. Each row represents each 50 years period, from top to bottom: mean productivity over the first 50 years (first line), over the 50 years after (second line) and over the last 50 years (third line). The table shows the productivity vulnerability (i.e. mean productivity is larger under climate change than under current climate) of each species for each site across three times.

#### 3.1.3 Stand vulnerability through stand basal area

In Bauges and Vercors, monospecific stands did not appear vulnerable, i.e. we found no difference between final basal area under current climate and under climate change, except for spruce in Vercors for which basal area declined by around 10%. In Ventoux, all monospecific stands were sharply vulnerable (Fig. 6), with a decrease between 48% (for fir stands) and 18% (for oak and spruce stands).

**Figure 5:**
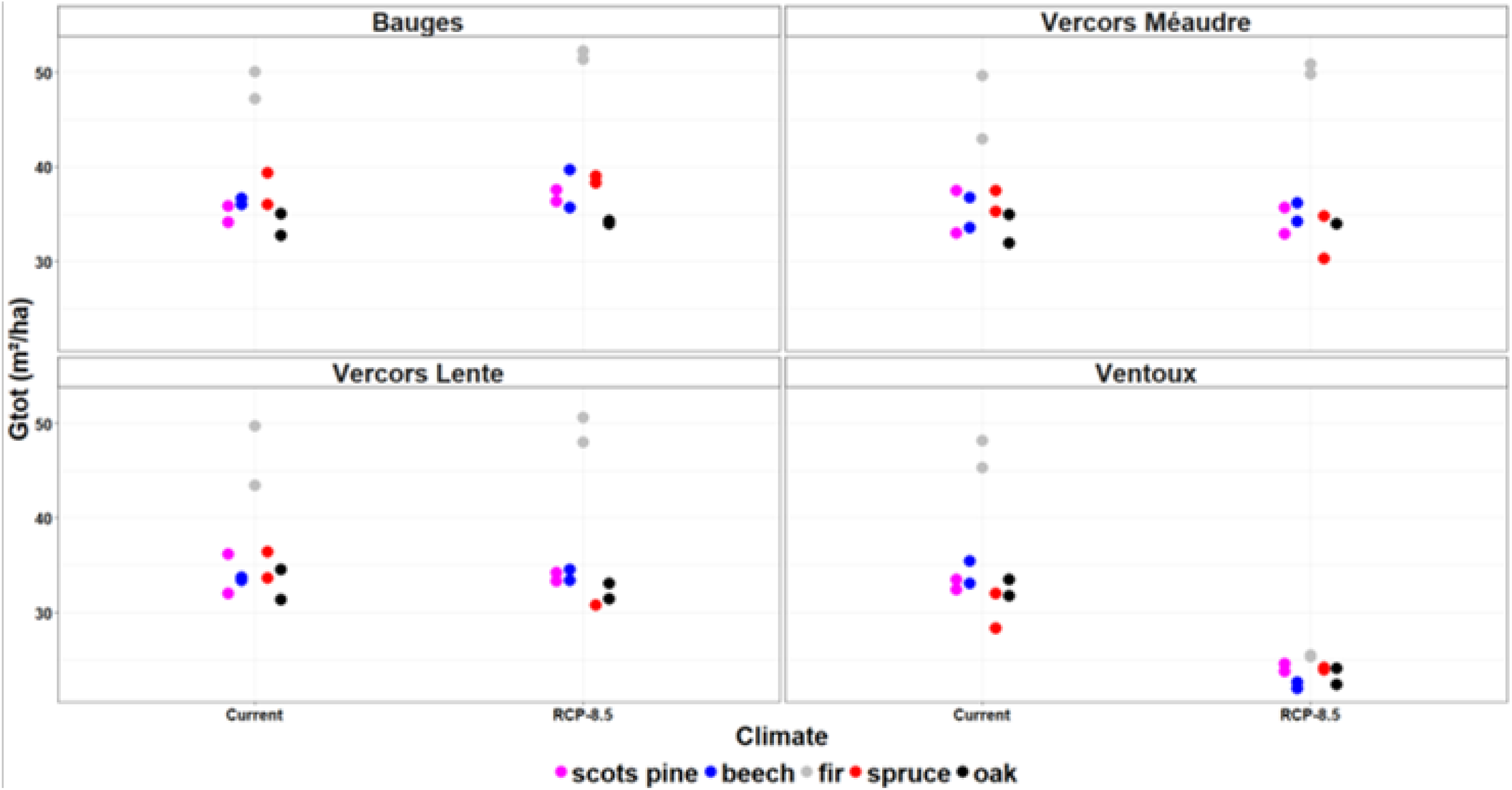
Total stand basal area after 150 years of the target species in monospecific stands without management for each set of climate conditions: current climate or climate change (RCP 8.5). Each species is represented separately: beech (blue), Scots pine (pink), fir (grey), spruce (red) and pubescent oak (black). Each panel corresponds to one site: Bauges, Ventoux, Vercors Lente and Vercors Méaudre.

**Figure 6:**
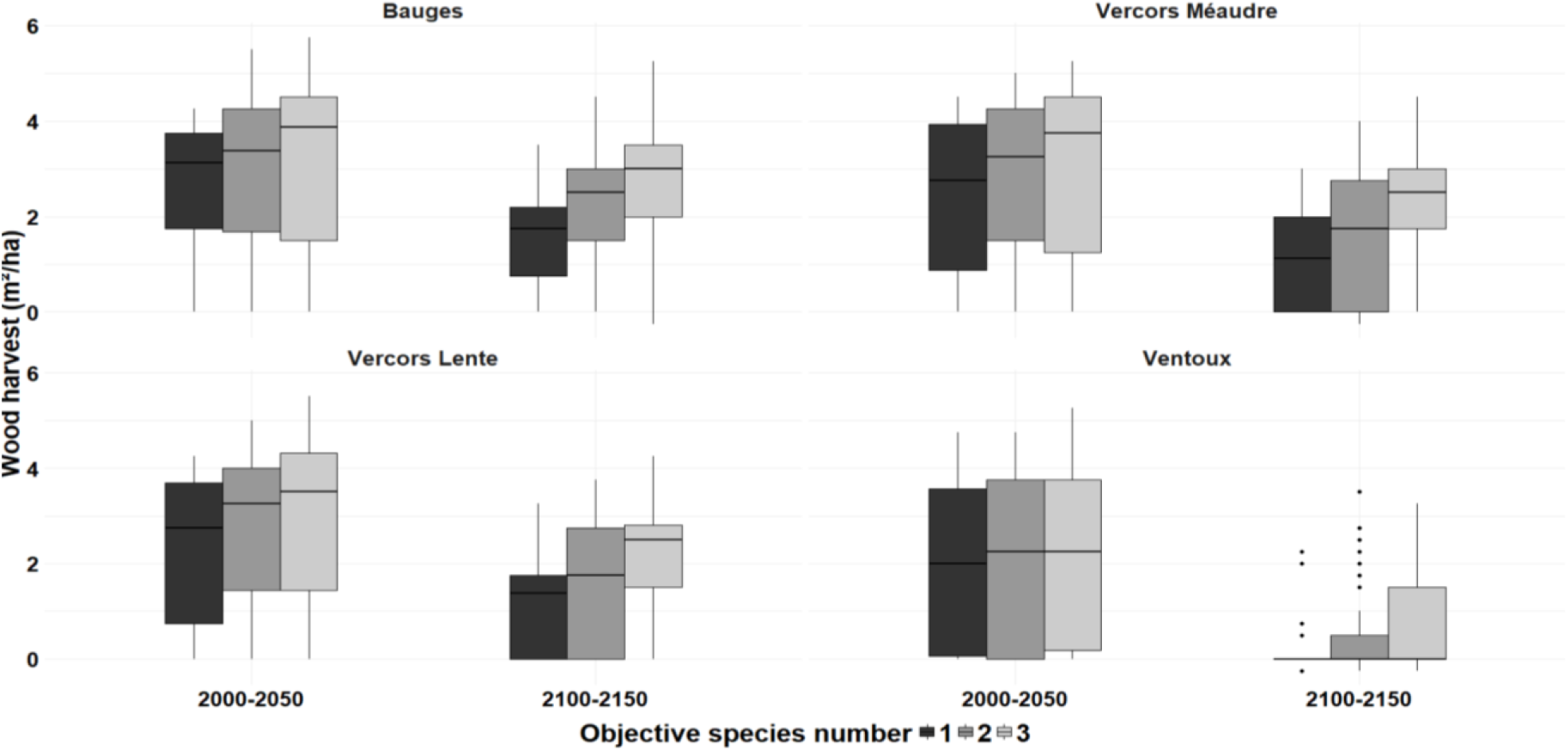
Wood harvest for initial (the first 50 years of the simulation) and final (the last 50 years of the simulation) time periods, with “stable management” scenarios. The management selected aims to stabilize initial stand composition. Species richness is represented by grey panel. Each part corresponds to one site: Bauges, Ventoux, Vercors Lente and Vercors Méaudre.

### 3.2 Would mixed stands or conversion stands mitigate climate change effects on forest?

#### 3.2.1 “Stable management” scenario

For all sites, initial stand species richness affected wood harvest. Wood harvest was not significantly different between monospecific and 2-species and 3-species mixed stands in the first 50 years (Fig. 7 and Table 1). In the last 50 years of 150 years, wood harvest was significantly different between monospecific and both 2- and 3-species mixed stands, but also between 2- and 3-species mixed stands, with wood harvest increasing with stand initial species richness, except in Ventoux (no difference between 2- and 3-species mixed stands) and Bauges (no difference between monospecific and 2-species mixed stands).

**Figure 7:**
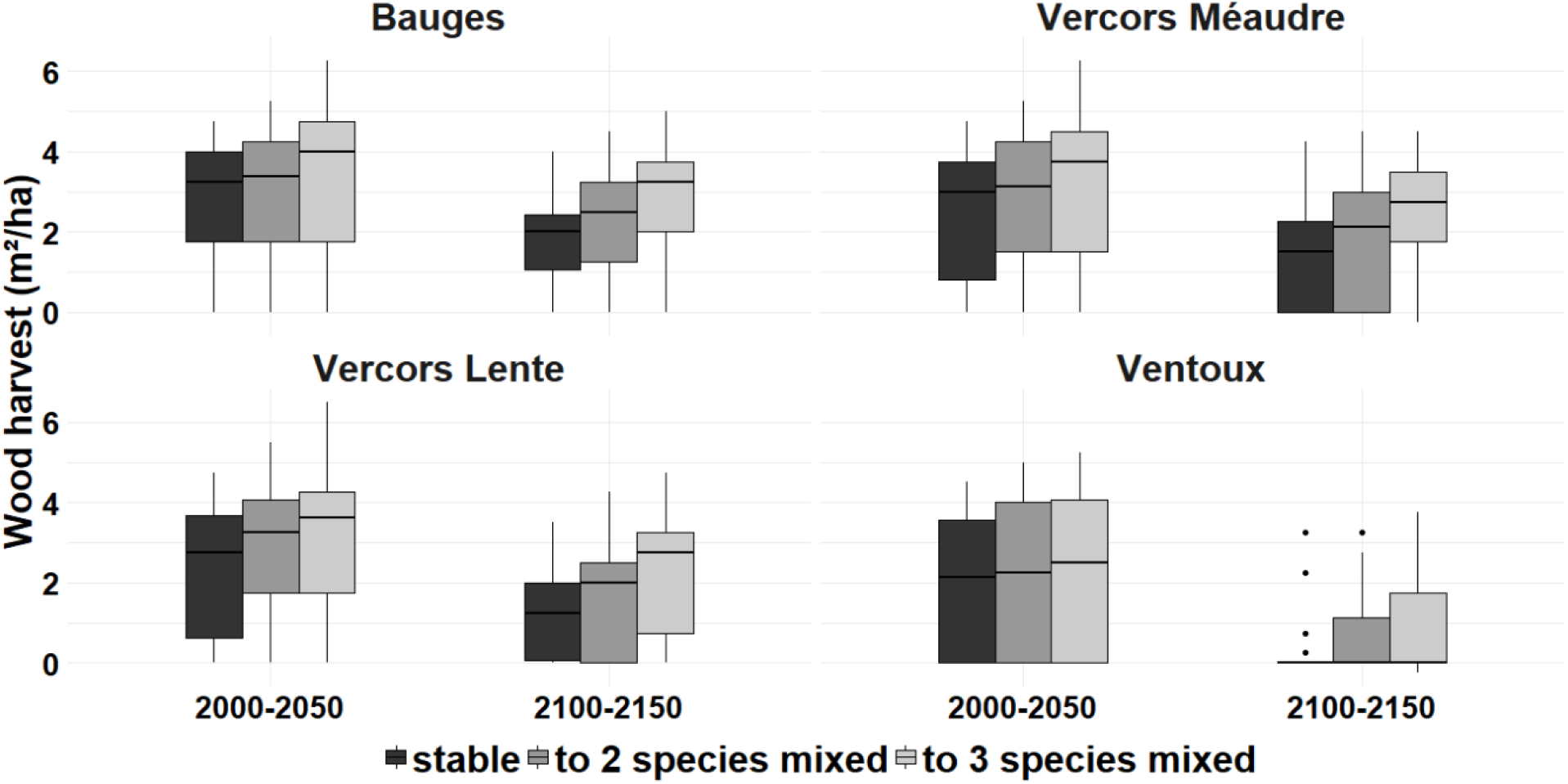
Wood harvest for initial (the first 50 years of the simulation) and final (the last 50 years of the simulation) time periods, with “conversion management” scenarios. The management selected aims to stabilize initial stand composition. Species richness is represented by grey panel. Each part corresponds to one site: Bauges, Ventoux, Vercors Lente and Vercors Méaudre.

**Table 1:** This table represents the p-value (student test) of mean harvest timber stand comparison, subject to stable management. Two periods are analyzed: the first 50 years and the last 50 years. We compared monospecific stands *vs.* 2-species mixed stands (1^st^ line) and *vs.* 3-species mixed stands (2^nd^ line), and 2- *vs.* 3-species mixed stands (3^rd^ line). In bold are represented significant difference (p-value < 0.05).

| Stand type | Time period | Bauges | Vercors Méaudre | Vercors Lente | Ventoux |
| --- | --- | --- | --- | --- | --- |
| monospecific *vs.* 2-species | 0-50 | 0.38 | 0.29 | 0.22 | 0.95 |
|  | 100-150 | 0.01 | **0.01** | **0.03** | 0.25 |
| monospecific *vs.* 3-species | 0-50 | 0.07 | 0.07 | 0.07 | 0.73 |
|  | 100-150 | **0.00** | **0.00** | **0.00** | **0.04** |
| 2-species *vs.* 3-species | 0-50 | 0.11 | 0.16 | 0.25 | 0.44 |
|  | 100-150 | **0.00** | **0.00** | **0.00** | 0.14 |

#### 3.2.2 “Conversion management” scenario

Wood harvest was correlated with stand conversion. For all sites, wood harvest at long term, i.e. between 100 and 150 years, depended strongly on stand targeted species richness. In last 50 years, wood harvest of monospecific stands managed in conversion to 3-species mixed stands was significantly higher than for monospecific stands with “stable” management (Fig. 8 and Table 2), except in Ventoux.

**Table 2:** This table represents the p-value (student test) of mean harvest timber stand comparison, subject to conversion management. Two periods are analyzed: the first 50 years and the last 50 years. We compare monospecific stand with 2-species mixed stand (1st line) and with 3-species mixed stand (2nd line), and 2 and 3-species mixed stand (3rd line). In bold are represented significant difference (p-value < 0.05).

| Stand type | Period | Bauges | Vercors<br>Méaudre | Vercors<br>Lente | Ventoux |
| --- | --- | --- | --- | --- | --- |
| monospecific vs 2-species | 0-50 | 0.47 | 0.29 | 0.33 | 0.60 |
|  | 100-150 | 0.13 | 0.12 | 0.11 | 0.28 |
| monospecific vs 3-species | 0-50 | 0.07 | **0.04** | 0.06 | 0.31 |
|  | 100-150 | **0.00** | **0.00** | **0.00** | **0.03** |
| 2-species vs 3-species | 0-50 | 0.08 | 0.12 | 0.14 | 0.44 |
|  | 100-150 | **0.00** | **0.00** | **0.00** | 0.08 |

Thus, increasing species richness up to 3 species increased the possible wood harvest, except in the Ventoux site (Fig. 8 and Table 2). Therefore, if maximizing wood harvest was a key objective, then managing forests as mixed stands management appeared as a relevant option under climate change, at least in French Alps.

We also observed that stands experiencing the conversion management allowed to reach similar levels of harvested wood after 150 years than stands managed as mixed stands since the beginning of the simulation. Moreover, increasing diversity effect had the same pattern for both management scenarios. Moreover, conversion and stable mixed-stand managements were equivalent in wood harvested, but depending on species, site and period considered (Table S4). Conversion management of monospecific stands induced different patterns depending of species: higher wood harvested than corresponding mixed stands (for fir), lower wood harvested (for pubescent oak and Scots pine, except in Ventoux), while for spruce and beech the pattern depended on site and period.

With a linear model (Table and Fig. S6), we concluded mixture management has a significant and positive effect on mean wood harvest. Moreover, increasing drought induced decreasing wood harvest. However, this trend became weaker with increasing species richness (for OS30). This result showed a buffer effect of mixture management on drought in our specific case.

## 4 DISCUSSION

### 4.1 Species vulnerability under climate change

Our simulations suggested that climate change may strongly alter monospecific stands functioning in our study area (French Alps), depending on site and species considered. This pattern echoes recent projections for the French Alps (Mina et al., 2017) indicating that climate change induces large alterations in the supply of several ecosystem services (timber production, carbon storage, protection against rockfall and avalanches) and biodiversity conservation.

In this study, we considered the three following situations (considering current climate) about the studied species: species at their range limit, weak competitive species in middle of their range, and strong competitive species in the middle of their range. In our study, the first situation corresponds to fir and beech stands in Ventoux and spruce stands in Vercors. The second situation corresponds to oak in all sites and Scots pine in Ventoux. The third situation corresponds to spruce in Bauges, and fir and beech in Bauges and Vercors. As Spruce was not adapted in Ventoux in current climate, we did not discuss this case further.

Monospecific fir and beech stands in Ventoux are highly vulnerable to dry conditions and soil water deficit as summarized in Bréda et al. (2006b) review (for fir see also Lebourgeois et al., 2013; Mina et al., 2015; Sánchez-Salguero et al., 2017). The trend depicted by our results trend consistent with this review: basal area of target species decreased very quickly (barely 50 years after the beginning of the simulation) and dropped to 10% of total proportion after 150 years under climate change. The gaps in canopy induced by mortality events allowed pioneer or post-pioneer species (e.g. Scots pine or oak) to colonize the patch (after 100 years).. Thus, the mean productivity of monospecific stands did not seem vulnerable after 150 years in Ventoux because Scots pine and oak have replaced the strong competitor but much less tolerant to drought species (either spruce, fir or beech). However, some studies found higher vulnerability of Scots pine at its southern range limits in Switzerland (Bigler et al., 2006b; Dobbertin et al., 2005), which may reduce competitive advantage of Scots pine in case of severe climate change. Moreover, pubescent oak and Scots pine seems to be sensitive to soil and atmospheric water deficits (Poyatos et al., 2008), which could also impact mean productivity under repeated drought events.

For spruce stands in Vercors (in the Lente and Méaudre sites), the same trend was found. Spruce proportion decreased because of drier and warmer conditions. High spruce vulnerability was already identified in Alps thank to observed data in past and current climate (Hartl-Meier et al., 2014; Levesque et al., 2013), because of a high sensitivity to drought (see Lu et al., 1996 in Vosges mountain). Oak could benefit from the disappearance of spruce. However, decreasing proportion of most drought sensitive species induced increasing proportion of most tolerant species (Niinemets and Valladares, 2006) that were subjected to strong competitive interactions in current climate.

### 4.2 Effect of mixed management on stands productivity

In our study, mixed-stand management seemed to strongly increase wood harvest after 100 years of simulation without increasing the vulnerability of the stands, allowing to harvest wood without negative impact on forest ecosystem. Our conclusion highlight mixed management effect on forest productivity and not mixed stand effect on productivity, but this does not exclude that mixed stands without management are also less vulnerable than monospecific stand.

According to previous studies, mixed stand management was put forward as a good option to maintain forest functioning and services (e.g.spruce-birch stand, Felton et al 2010) accelerating conversion from monospecific to mixed stand or maintaining mixed stand. Mixed stands have actually been reported to reduce species sensitivity to drought (fir in fir-beech or fir-spruce stands, Lebourgeois et al., 2013, but see also Grossiord et al., 2014b) and/or increase (beech in beech-oak stands, Pretzsch et al., 2013; beech-spruce stand, Pretzsch et al., 2014; or Scots pine-oak stand, Steckel et al., 2019) or stabilize (beech-Scots pine stand, Del Río et al., 2017; beech-fir stands, Jourdan et al., 2019) species productivity when compared to monospecific stands.

In our simulations, monospecific stands of species at their range limit cannot be maintained as monospecific stands in the coming decades (spruce, fir and beech in Ventoux, and spruce in Vercors), because aimed species were too vulnerable to climate change. This is for instance in agreement with Mina et al.' simulation study (2017) that found a sharp decrease of spruce proportion in mixed Vercors' stand under severe climate change scenarios and under different management scenarios. Moreover, Hlásny et al. (2017) showed that managing forests with higher diversity induces higher wood production of spruce in Eastern Alps with climate change (i.e. with higher temperature increase) compared to monospecific stand management.

In Ventoux, there is complete initial species substitution of late- or middle-succession species, i.e. spruce, fir and beech, by early-succession species, i.e. Scots pine and oak. Such substitution pattern has been predicted with correlative species distribution models in North American forest ecosystems (Iverson and Prasad, 2001) and in Mediterranean-alpine ecosystems (Benito et al., 2011). This trend is perhaps misestimated because we considered a limited number of competitive species under stressful conditions (Scots pine and pubescent oak). However, one should notice that other early-succession species (other pine species, for example) may change Scots pine or pubescent oak mean productivity (Riofrío et al., 2017).

### 4.3 Inputs of ForCEEPS

In most forest-oriented studies, climate change is mimicked *via* a latitudinal gradient of sites in empirical studies (e.g. in the Alps, Pretzsch et al., 2010) or *via* rainfall exclusion in experimental studies (Estiarte et al., 2016). Regarding empirical studies, such an approach relies on the “space for time substitution” to assess climate change impacts on ecosystems (Blois et al., 2013), which is an efficient way to assess climate change impacts of forest stands. However, such field-based approaches relying on gradients allow a limited climatically analogy (Vallet and Perot, 2018). Moreover, testing forest management scenarios *in situ* is a difficult task, with long-term studies (Gavinet et al., 2019). Therefore, additionally considering the impact of climate change with a latitudinal or altitudinal gradient accentuated the difficulties. Modelling approaches make it possible to simulate the dynamics of forest stands, while considering climate change and management. With simulation-based studies, it is possible to have a comprehensive view of the stand development over time and on processes driving observed patterns, in the range of applicability of the model of course. More generally, to deal with the high uncertainty characterizing future climate, forest models appear as a key tool to test various scenarios (Cordonnier et al., 2018, Morin et al., 2020), complementing empirical and experimental approaches.

However, it must be reminded that the simulations relied on simplified mechanisms compared to real processes and stochastic events involved in ecosystem functioning. For example, competitive dynamics in ForCEEPS are focused on light acquisition and do not explicitly consider tree roots. Thus, competition for water and nutriments acquisition is only indirectly considered. Moreover, other uncertainties are due to lack of knowledge on forest dynamics under climate change, but also difficulties to reproduce some critical mechanisms (like photosynthesis or respiration outside the past temperature range). Other modelling approaches could have been used, like process-based models in which productivity is simulated through the outcome of respiration and photosynthesis processes of the different compartments of the forest ecosystem (Dufrêne et al., 2005; Jonard et al., 2020; Landsberg et al., 2003). However, ForCEEPS presents a balance between complexity and generality, and is notably easily calibrated for new species as it requires only a limited number of parameters (Morin et al., 2020a). Furthermore, a perspective to obtain more robust simulations could be to couple ForCEEPS with a process-based model.

Nevertheless, while keeping in mind its limits, the model ForCEEPS has a great potential to be a relevant tool in both functional ecology and forest management, notably because it is relatively easy to calibrate for many species. Regarding functional ecology, using such a model would for instance allow testing hypotheses related to changing interspecific dominance under climate change. Regarding forest management, such a model could allow to test forestry itineraries under various climate scenarios, which very few models can achieve so far.

### 4.4 Perspectives for management

In this study, we explored how management may help in maintaining forest cover and functioning to preserve and maybe even improve some ecosystem services provided by forests. Because our results rely on simulations, our discussion on management should not be taken as recommendation, but as open debate. Furthermore, we focus here on wood harvest and forest cover, but obviously management decisions must consider as many ecosystem services as van der Plas et al. (2016).

#### 4.4.1 Mixed stand management

Our simulations showed that mixed stands might be promoted with three species mixtures. Our results also strongly suggest that converting monospecific stands may significantly improve stand performance in terms of wood harvest mean and decrease the vulnerability to climate change (see Table S5). Such a result could come from both a selection effect and a complementary effect (Loreau and Hector, 2001), and our experiment cannot disentangle these two mechanisms properly. It is thus possible that one adapted species could lead to higher productivity with monospecific stand management (for example, pubescent oak in Ventoux) compare to mixed stands management. Nevertheless, the knowledge on which composition, including monospecific stands, performs best would provide more information to managers on possible management options to ensure the production function in the face of climate change.

One interesting conclusion of our study is that the targeted composition was not necessarily the final composition, indicating that some mixed stands are more difficult to manage than others (Bauhus et al., 2017; Cordonnier et al., 2018). For example, in “stable management”, a significant part of simulations did not reach the targeted species composition after 150 years (Fig. S6). This shows that instead of fixing an *a priori* given composition, managers could instead adopt a management approach that accompanies the natural dynamics of mixtures provided that the new species are adapted to the anticipated future climate. This approach would better fit the idea of adaptive management (Rist et al., 2013) that takes advantage of unpredicted events.

#### 4.4.2 For thought about forest management in context of climate change

Some studies recommended to use interventionist silvicultural practice, promoting nonnative species and non-local provenances (in European forest, Brang et al., 2014 or Canadian forest Leech et al., 2011) or genetic engineering (Dumroese et al., 2015). Although this type of management is very controversial (Aubin et al., 2011; Pedlar et al., 2012; Winder et al., 2011), in mountain forest context, it could be invoked to maintain forest coverage, to protect human population against erosion, rock fall or avalanche. In addition, recent studies have shown the advantage of keeping a continuous forest cover in order to maintain microclimate forest buffering warming (Zellweger et al., 2020). One should recall here that our study focused on managed forests to compare mixed and monospecific forests. Our results only relate to this comparison. Thus, we cannot drive any conclusion about the effect of management on forests' vulnerability to climate change.

In our study, despite temperature and precipitation changes, pubescent oak seemed to remain less competitive than fir and beech in Vercors. This would suggest that favoring a shift in composition towards less drought-sensitive species in these sites – for instance with assisted migration (McLachlan et al., 2007; Vitt et al., 2010) – does not always appear relevant, while other management actions, like specific “favorable” forestry, could be sufficient. “Favorable” forestry for one species corresponds to the promotion of the development of this species through stand management. Contrariwise, if forest cover decreases drastically, assisted migration of Mediterranean species *(Quercus pubescens, Quercus ilex, Pinus halepensis or Pinus pinea)* could become an interesting option to maintain forest cover and limit soil erosion. In our case one type of migration could be relevant: translocation just beyond the range limit (assisted range expansion; Leech et al. 2011). Indeed limited dispersal abilities and/or highly fragmented habitat (Vitt et al., 2010) can induce difficulties for some species to colonize available nearby habitat. In theory, physical factors (soil types, topography, photoperiod) and biotic factors (community composition) of recipient forests are very close to the native range of the species and may, correspond to the previous range of the species on longer time scales (Hoegh-Guldberg et al., 2008; Hunter, 2007). In case of Scots pine, which is already widely present in South of Alps -more than half 2018-IFN inventory contained Scots pine trees (51%)-, this strategy of management seems appropriate.

### 4.5 Research perspectives

We tested a limited number of combinations of “species composition-climate scenario-management scenario”. Therefore, these should rather be considered as trends depicted at the forest level and not to precise recommendations at the stand level. New simulations should be added to follow up the present work and determine more precisely which management would allow optimal wood harvest, for each site. For instance, it might be interesting to test mixtures with more than 3 species. Positive diversity effects on average productivity and its stability (Jucker et al., 2016, 2014b) have been shown in observation-based studies, but these studies remain rare and do not allow to study the same species and the same species composition on an extended latitudinal gradient. Our modelling approach could easily study the same species over latitudinal gradients by controlling species proportion with 4 or 5 species. For instance, we could add other species, especially for Ventoux, such as other pine species (*Pinus pinaster, Pinus nigra or Pinus halepensis,* for wood industry and *Pinus pinea,* for non-wood forest product), other oaks species *(Quercus ilex, Quercus pyrenaica or Quercus suber*, for example) or even non-native species (*Robinia pseudoacacia*).

Regarding the model outputs, it would be highly relevant to extend the range of ecosystem services studied. We briefly mentioned the protective forest role in mountain areas and conservation-related services, but there are many other services that may considered, if the model allows it: maintenance of micro-habitat (Courbaud et al., 2016), maintenance of carbon and nutrients cycles functioning (Corbeels et al., 2005), carbon storage (Delpierre et al., 2012) or spiritual or recreational dimension (McFarlane and Boxal, 2000). A natural extension of the present study could also focus on carbon sequestration, by quantifying carbon stored in standing trees and carbon stocked in wood products. Finally, considering operational cost of management actions could also be included in future studies. This constraint is very strong on French alpine productive forests and must be fully integrated. More generally, it would be crucial to consider other ecosystems services as important to maintain as wood production, to test various scenarios related to different forest policies.

## AUTHORS’ CONTRIBUTIONS

XM, CR, PD, TC and MJ designed the research and methodology. XM and FdC developed the model, parameterized the species, including the management module, with the help of PD. MJ and BC carried-out the simulations; MJ analyzed the data; MJ and XM led the writing of the manuscript. All authors contributed critically to the drafts and gave final approval for publication.

## ACKNOWLEDGMENTS

MJ benefited from an ADEME grant. This study was mostly funded by the project DISTIMACC (ECOFOR-2014-23, French Ministry of Ecology and Sustainable Development, French Ministry of Agriculture and Forest) and benefited from the ANR project BioProFor (contract no. 11-PDOC-030-01). This study strongly benefited from the help of B. Cornet, for his assistance in pre-analyses. We thank F. de Coligny for his help on the development of the ForCEEPS model. This work was based on the Capsis platform: http://www.inra.fr/capsis.

## Notes

### Competing Interest Statement

The authors have declared no competing interest.

