## Supplementary Materials for "Managing mixed stands can mitigate severe climate change impacts on ecosystem functioning"

#### Appendix 1: *Outline of the article*

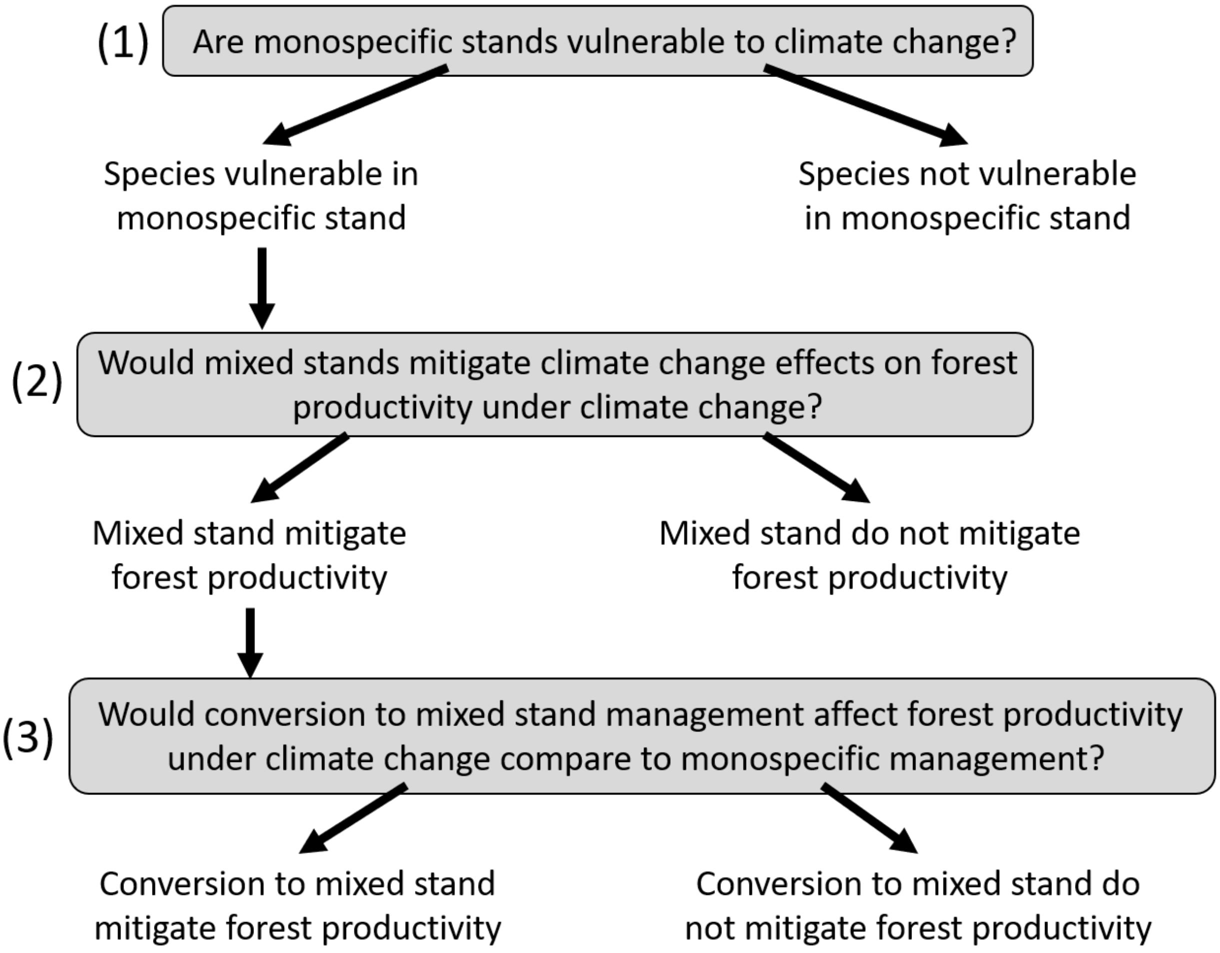

### Appendix 2: *Description of ForCEEPS management module*

The management module is optional in ForCEEPS. It allows to consider logging effect on forest dynamic, reproducing forest thinning. To reproduce as best as possible current management, four options are implemented (see below): thinning frequency and intensity, aimed proportion and thinning type.

Thinning frequency and intensity can be initializing for each simulation. In our study, the frequency is 12 years (according to foresters' practices in these sites, ONF comm.). This does not depend on stand variables (i.e. basal area, volume, species composition) and does not change during the simulation. Thinning intensity can be set as a percentage of basal area remaining after thinning or as an objective basal area after thinning (also reproducing French foresters' practices, ONF comm.). During thinning, trees are removed randomly or according thinning rules, until targeted basal area or targeted percentage of basal area is reached.

The type of thinning type can be initialized. It is possible to choose to set a targeted proportion for each species for each cutting operation. It is also possible to choose thinning from top (largest diameters are privileged during thinning) or from bottom (thinnest diameters are privileged during thinning). In our study we choose to use top-thinning, which closer to French sylvicultural practices.

Note that in the present study we also did not impose a minimum threshold of the thinning operation (in terms of basal area), although the model allows it.

### Appendix 3: *Site characteristics*

| Site | Bauges | | Vercors Méaudre | | Vercors Lente | | Ventoux | |
|---|---|---|---|---|---|---|---|---|
| | low | high | low | high | low | high | low | high |
| latitude | 45.697930°N/ 6.214553°E | | - | | 44.928504°N / 5.321516°E | | 44.187901°N / 5.253608°E | |
| *Current climate* | | | | | | | | |
| • Temperature (°C) | 7.7 | 6.5 | 7.7 | 5.1 | 7.3 | 5.3 | 8.3 | 6.2 |
| • Precipitation (mm) | 1770 | 1770 | 1301 | 1301 | 1346 | 1346 | 992 | 992 |
| *Climate change* | | | | | | | | |
| • Temperature (°C) | 11.7 | 10.5 | 11.7 | 9.1 | 11.1 | 9.1 | 12.1 | 12.1 |
| • Precipitation (mm) | 1833 | 1833 | 1352 | 1352 | 1197 | 1197 | 882 | 882 |
| *Site characteristic* | | | | | | | | |
| • Elevation (m) | 1000 | 1300 | 1000 | 1300 | 1000 | 1300 | 1000 | 1300 |
| • Bucket size | 50 | | 30 | | 30 | | 25 | |
| • Max ETP rate | 12 | | 12 | | 12 | | 12 | |
| • Nitrogen | 100 | | 100 | | 100 | | 100 | |

Table summarizing site characteristics used for ForCEEPS. Bauges, Vercors Méaudre, Vercors Lente and Ventoux are characterized by coordinates, current climate and climate change variables (temperature and precipitation) and site characteristics (elevation, bucket size, max EPT rate and soil nitrogen).

### Appendix 4: *Targeted species proportion*

| **Site** | **Bauges** | | **Vercors Méaudre** | | **Vercors Lente** | | **Ventoux** | |
|---|---|---|---|---|---|---|---|---|
| **Climate** | **Current** | **RCP 8.5** | **Current** | **RCP 8.5** | **Current** | **RCP 8.5** | **Current** | **RCP 8.5** |
| **Spruce** | 0.68 | 0.44 | 0.72 | 0.16 | 0.73 | 0.09 | 0.11 | 0 |
| **Fir** | 0.65 | 0.57 | 0.63 | 0.60 | 0.64 | 0.6 | 0.72 | 0 |
| **Beech** | 0.75 | 0.58 | 0.71 | 0.62 | 0.73 | 0.61 | 0.78 | 0.03 |
| **Scots pine** | 0.16 | 0.09 | 0.20 | 0.10 | 0.17 | 0.09 | 0.22 | 0.55 |
| **Oak** | 0.61 | 0.62 | 0.48 | 0.67 | 0.41 | 0.65 | 0.63 | 0.95 |

Table summarizing targeted species proportion after 150 years without management (*spruce, fir, beech, scotes pine, oak*) in each site (*Bauges, Vercors Méaudre, Vercors Lente, Ventoux*) and for each climate (*current* and *RCP8.5*)

### Appendix 5: *Table of wood harvested for mixed management*

| Diversity | species | site | 2000-2050 | | | 2100-2150 | | |
|---|---|---|---|---|---|---|---|---|
| | | | Stable (50-50) | conversion | | Stable (50-50) | conversion | |
| 2-species | Beech | Bauges | 2.91 | 2.72 | S | 2.30 | 2.35 | C |
| | | Ventoux | 2.29 | 2.56 | C | 0.27 | 0.31 | C |
| | | Vercors_L | 2.7 | 2.61 | S | 1.92 | 1.85 | S |
| | | Vercors_M | 2.75 | 2.58 | S | 1.95 | 2.03 | C |
| | Spruce | Bauges | 3.24 | 3.30 | C | 2.44 | 2.60 | C |
| | | Ventoux | 1.10 | 0.56 | S | 0.38 | 0.35 | = |
| | | Vercors_L | 3.05 | 2.93 | S | 1.24 | 0.90 | S |
| | | Vercors_M | 3.09 | 2.95 | S | 1.64 | 1.32 | S |
| | Fir | Bauges | 3.5 | 4 | C | 2.69 | 2.92 | C |
| | | Ventoux | 2.48 | 3.46 | C | 0.32 | 0.41 | C |
| | | Vercors_L | 3.09 | 3.84 | C | 2.42 | 2.64 | C |
| | | Vercors_M | 3.2 | 3.91 | C | 2.25 | 2.68 | C |
| | Scots pine | Bauges | 2.68 | 2.38 | S | 1.80 | 1.77 | S |
| | | Ventoux | 2.02 | 2.25 | C | 0.72 | 0.95 | C |
| | | Vercors_L | 2.38 | 2.39 | = | 1.42 | 1.44 | = |
| | | Vercors_M | 2.42 | 2.30 | S | 1.55 | 1.49 | S |
| | Pubescent oak | Bauges | 2.98 | 2.21 | S | 2.21 | 1.52 | S |
| | | Ventoux | 1.99 | 2.18 | C | 0.48 | 0.79 | C |
| | | Vercors_L | 2.74 | 1.97 | S | 1.68 | 1.14 | S |
| | | Vercors_M | 2.78 | 2.19 | S | 1.80 | 1.33 | S |
| 3-species | Beech | Bauges | 3.15 | 2.94 | S | 2.89 | 3.06 | C |
| | | Ventoux | 2.3 | 2.64 | C | 0.59 | 0.60 | = |
| | | Vercors_L | 2.88 | 2.73 | S | 2.15 | 2.27 | C |
| | | Vercors_M | 2.96 | 2.69 | S | 2.29 | 2.59 | C |
| | Spruce | Bauges | 3.35 | 3.51 | C | 2.95 | 2.99 | C |
| | | Ventoux | 1.78 | 0.71 | S | 0.56 | 0.62 | C |
| | | Vercors_L | 3.13 | 3.25 | C | 1.87 | 1.99 | C |
| | | Vercors_M | 3.22 | 3.32 | C | 2.09 | 2.12 | C |
| | Fir | Bauges | 3.41 | 4.63 | C | 3.06 | 3.34 | C |
| | | Ventoux | 2.33 | 3.79 | C | 0.52 | 0.70 | C |
| | | Vercors_L | 3.11 | 4.35 | C | 2.38 | 2.94 | C |
| | | Vercors_M | 3.17 | 4.46 | C | 2.45 | 3.01 | C |
| | Scots pine | Bauges | 3.03 | 2.81 | S | 2.46 | 2.15 | S |
| | | Ventoux | 2.12 | 2.37 | C | 0.82 | 1.01 | C |
| | | Vercors_L | 2.77 | 2.69 | S | 1.91 | 1.86 | S |
| | | Vercors_M | 2.85 | 2.63 | S | 2.01 | 2.01 | = |
| | Pubescent oak | Bauges | 3.18 | 2.48 | S | 2.78 | 2.38 | S |
| | | Ventoux | 2.14 | 2.31 | C | 0.65 | 0.86 | C |
| | | Vercors_L | 2.92 | 2.17 | S | 2.04 | 1.92 | S |
| | | Vercors_M | 2.99 | 2.42 | S | 2.17 | 1.99 | S |

Comparison of wood harvested in 2 period (2000-2050 and 2100-2150) between Stable management of 2 species (50%-50%) or 3 species (40%-30%-30%) mixed stand and conversion management of monospecific stand to 2 or 3 species. Each species (beech, fir, spruce, Scots pine, Pubescent oak) and site (Bauges, Vercors and Ventoux) are considered separately.

### Appendix 6: *Linear model explaining wood harvest average*

| **Model: Pm_50_3 ~ Precipitation + diversity + Precipitation:diversity + cst** | | | | | | **$R^2$** |
|---|---|---|---|---|---|---|
| | | Estimate(±SD) | Estimate(±SD) | Estimate(±SD) | Estimate(±SD) | |
| **OS20** | | **0.002 (±0.0003)** | **0.57 (±0.20)** | -0.0001 (±0.0001) | **-1.96 (±0.32)** | **0.44** |
| **OS30** | | **0.002 (±0.0002)** | **0.43 (±0.13)** | **-0.0005 (±0.0001)** | **-2.04 (±0.24)** | **0.23** |
| **OS80** | | **0.001 (±0.0002)** | **0.52 (±0.15)** | -0.0000 (±0.0001) | -0.15 (±0.23) | **0.48** |

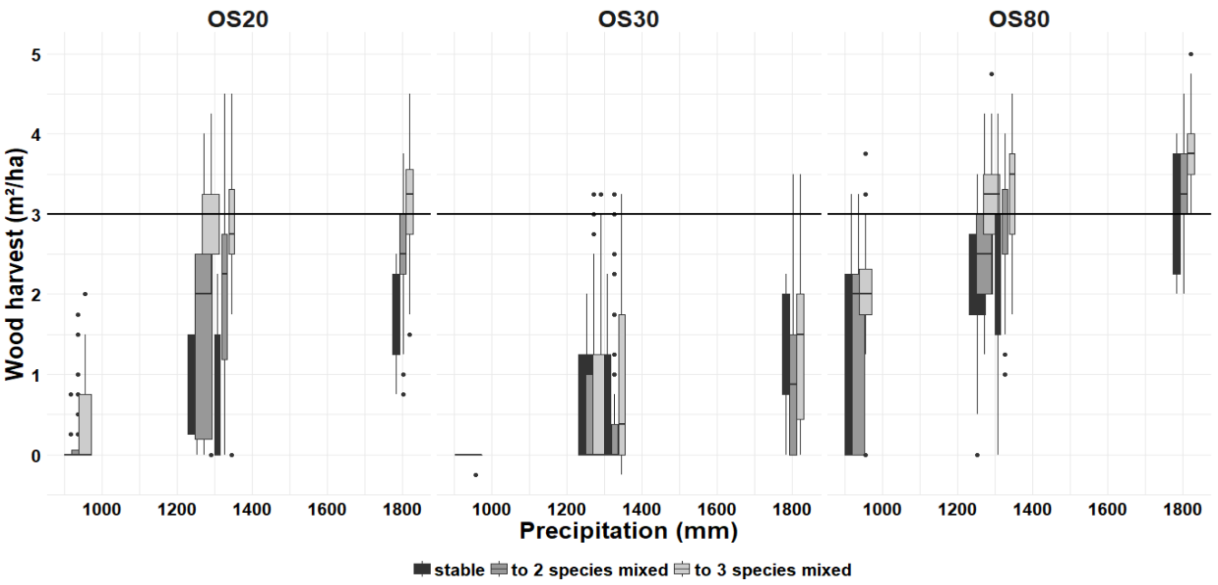


This appendix represents linear model explaining wood harvest average between 100 and 150 years of simulation with conversion management in function of: precipitation, diversity and the interaction, with summary table and corresponding figures. We fit one model for each basal area objective scenarios -20m²/ha (OS20), 30m²/ha (OS30) and 80% of previous basal area (OS80)-, because this variable influence greatly wood logging average. Table represents estimate (±standard deviation) of each explaining variable, bolded if p-value>0.05, and $R^2$ of each model. Figures represents wood harvest average in function of precipitation, considering separately each objective diversity.

### Appendix 7: *Real final proportion*

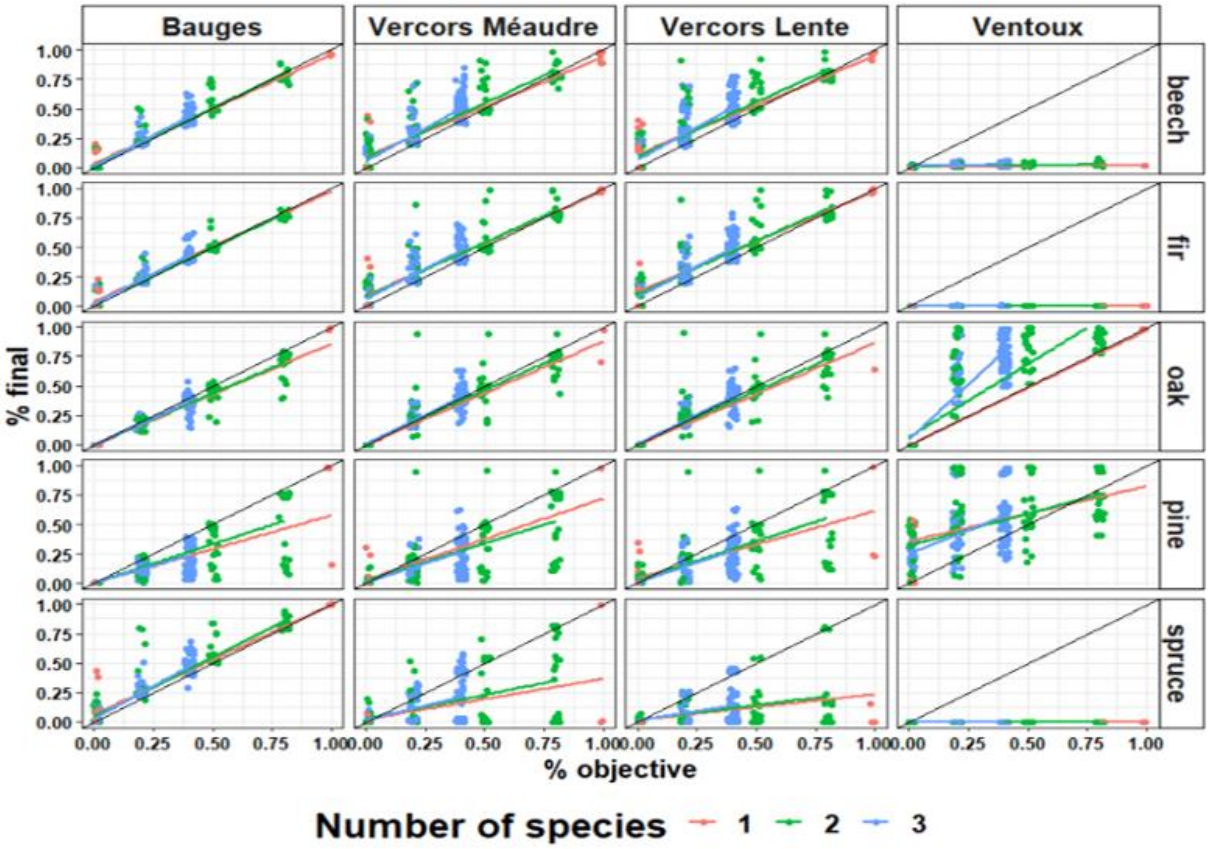


This graphic represents real final proportion in function of aimed proportion for each species (beech, fir, pubescent oak, scots pine and spruce). Each site is presented separately: Bauges, Vercors Méaudre, Vercors Lente and Ventoux. We also materialize levels of objective diversity with colors: monospecific stand (red), 2-species mixed stand (green) and 3-species mixed stand (blue). This figure allows to compare objective diversity and observed diversity in each stand.
